# c-Src induced vascular malformations require localised matrix degradation at focal adhesions

**DOI:** 10.1101/2023.09.15.557868

**Authors:** Patricia Essebier, Lilian Schimmel, Teodor Yordanov, Mikaela Keyser, Alexander Yu, Ivar Noordstra, Brittany Hill, Alpha S. Yap, Samantha J. Stehbens, Anne K. Lagendijk, Emma J. Gordon

## Abstract

Endothelial cells lining the blood vessel wall communicate intricately with the surrounding extracellular matrix, translating mechanical cues into biochemical signals. Moreover, vessels require the capability to enzymatically degrade the matrix surrounding them, to facilitate vascular expansion. c-Src plays a key role in blood vessel growth, with its loss in the endothelium reducing vessel sprouting and focal adhesion signalling. Here, we show that constitutive activation of c-Src in endothelial cells results in rapid vascular expansion, operating independently of growth factor stimulation or fluid shear stress forces. This is driven by an increase in focal adhesion signalling and size, with enhancement of localised secretion of matrix metalloproteinases responsible for extracellular matrix remodelling. Inhibition of matrix metalloproteinase activity results in a robust rescue of the vascular expansion elicited by heightened c-Src activity. This supports the premise that moderating focal adhesion-related events and matrix degradation can counteract abnormal vascular expansion, with implications for pathologies driven by unusual vascular morphologies.

## Introduction

The blood vascular network is largely formed by sprouting angiogenesis and is required to supply oxygen and nutrients to all tissues in the body [1]. Controlled sprouting and patterning of the vasculature is essential during embryonic development, with aberrant angiogenesis also occurring later in life in response to tissue injury or in disease. The binding of vascular endothelial growth factors (VEGFs) to vascular endothelial growth factor receptors (VEGFRs) is considered to be largely responsible for initiating sprouting [2], driving downstream signalling to induce endothelial cell (EC) identity, migration and proliferation. Sprouting angiogenesis and vascular integrity is mediated by endothelial cell-cell contacts, which in turn are controlled by the junctional localisation of Vascular Endothelial-cadherin (VE-cadherin) [3, 4]. Angiogenesis and cell-cell adhesion are also intricately associated with cell-matrix adhesions, with increasing evidence that if adhesions are disrupted, ECs display altered VE-cadherin localisation, sprouting and permeability [5–9].

Cell anchoring to the extracellular matrix (ECM) is primarily mediated through focal adhesions (FAs), which are large, dynamic, multi-protein complexes [10]. Matrix-bound transmembrane integrin receptors interact with intracellular signalling molecules, including focal adhesion kinase (FAK), c-Src and paxillin. Within focal adhesions, the mechano-effector proteins vinculin and talin directly interact with the actin cytoskeleton, undergoing confirmational changes in response to tension via a process known as mechanotransduction [11, 12]. There is increasing evidence that FAs and cell-cell adhesions share significant crosstalk via actin scaffolding processes [13, 14].

It is well appreciated that the non-receptor Src family kinases play critical roles in both cell-cell adhesion and cell-matrix adhesion, with their localisation and activity regulated in a context-dependent manner. VE-cadherin phosphorylation by Src family kinases (SFK) at distinct tyrosine sites within its intracellular tail mediates its localisation in endothelial cells, and controls vascular integrity and angiogenic sprouting [15–20]. While c-Src was traditionally thought to be primarily responsible for phosphorylation of VE-cadherin, recent advances in Cre-driven endothelial-specific mouse models identified that the SFK Yes, rather than c-Src, mediates VE-cadherin phosphorylation, turnover and junctional plasticity [18]. In agreement, endothelial-specific deletion of c-Src does not lead to gross defects in VE-cadherin internalisation. Rather, loss of c-Src leads to reduced retinal angiogenesis due to loss of FA assembly and cell-matrix adhesion, resulting in loss of sprout stability [21]. This suggests that SFKs play different roles in FA to cell-cell adhesion crosstalk, leading to context-dependent roles in vascular growth and function.

In addition to facilitating adhesion to the surrounding environment, FAs have been shown to play a role in remodelling of the ECM, acting as sites of localised exocytosis. We previously demonstrated that microtubules target and anchor at FA, facilitating delivery of exocytic vesicles [22]. In the context of migrating keratinocytes, this results in localised secretion of the matrix metalloproteinase (MMP) MT1MMP/MMP14, releasing adhesions from the surrounding matrix, and resulting in focal adhesion turnover. This directly associates FAs with the intracellular machinery regulating exocytosis, a fundamental process where molecules are transported to the cell surface, where they can be either released into the extracellular space or integrated into the plasma membrane [23]. Microtubules act as the major tracks for transport within cells, where vesicles are transported along these dynamic, tube-like structures by kinesin or dynein motors to the plasma membrane for exocytosis [24]. Exocytosis is critical for a range of cellular processes such as synaptic neurotransmission, paracrine signalling, or the release of hydrolytic enzymes, allowing the turnover of FAs which is critical for coordinated cell movement [25]. While MMPs are known to be critically important for a wide range of vascular processes, including angiogenesis, morphogenesis and wound repair [26, 27], precisely how alterations in MMP activity and localisation are regulated, and whether this is mediated at endothelial FAs, remains unclear.

Here, we sought to investigate the consequence of elevated c-Src activity on EC adhesions and vascular behaviour. We found that introducing a constitutively active (Y527F) c-Src mutation (c-Src-CA) into ECs in 3D vascular models resulted in a vast ballooning phenotype. These vessels lacked functional angiogenic sprouts and a continuous cellular lining of the vessels. The c-Src-CA mutation induced large FAs, elevated paxillin and VE-cadherin phosphorylation, and loss of cell-cell junctions. We found that local secretion of MMPs at these enlarged FAs resulted in degradation of ECM components, with broad spectrum pharmacological inhibition of MMPs rescuing the vascular ballooning induced by constitutively active c-Src. Thus, our work clearly shows the importance of tightly controlled FA formation and turnover, mediated by c-Src, in enabling functional angiogenesis.

## Results

### Elevated c-Src activity induces vascular malformations

Loss of endothelial c-Src results in decreased vascular stability due to loss of cell-matrix adhesion [21]. We further sought to determine how elevated c-Src activity would affect EC behaviour. c-Src activity in cells is normally regulated by a glycine/serine-rich flexible linker at the C-terminus [28] (Supp Fig 1A). We generated several c-Src-mScarlet (mSc) fusion proteins including wildtype (c-Src-WT-mSc), constitutively active (c-Src-CA-mSc) with a Y527F mutation preventing phosphorylation of the inhibitory Tyr527 [29], or dominant negative (c-Src-DN-mSc) with Y527F/K295R mutations, holding the protein in its open confirmation to bind target proteins, but with a dysfunctional kinase domain [30] (Supp Fig 1A-C). Expression of the c-Src-mScarlet fusion proteins in HUVECs using a lentiviral based system led to a significant increase in c-Src protein levels, which was not altered by either Y527 or Y527F/K295R mutations (Supp Fig 1D-E).

To assess how elevated c-Src activity altered vascular sprouting in 3D, we embedded microcarrier beads coated with control and c-Src mutant expressing ECs within fibrin gels [31]. After 7 days, c-Src-WT cells displayed a modest increase in the number of sprouts, although there was no significant increase in vascular area (Fig 1A-C). Introduction of the c-Src-CA mutation induced a drastic alteration in morphology, where instead of distinctive sprouts, severe vascular ballooning was observed (Fig 1A-D, Supp Movie 1). The c-Src-DN mutation completely abrogated this phenotype (Fig 1A-D), suggesting that this ballooning morphology was a result of increased c-Src kinase activity. Additionally, the c-Src-DN cells displayed decreased vascular area and fewer sprouts than Ctrl or c-Src-WT cells (Fig 1B-D), confirming this mutation suppresses endogenous c-Src signalling in a dominant negative fashion.

**Figure 1.**
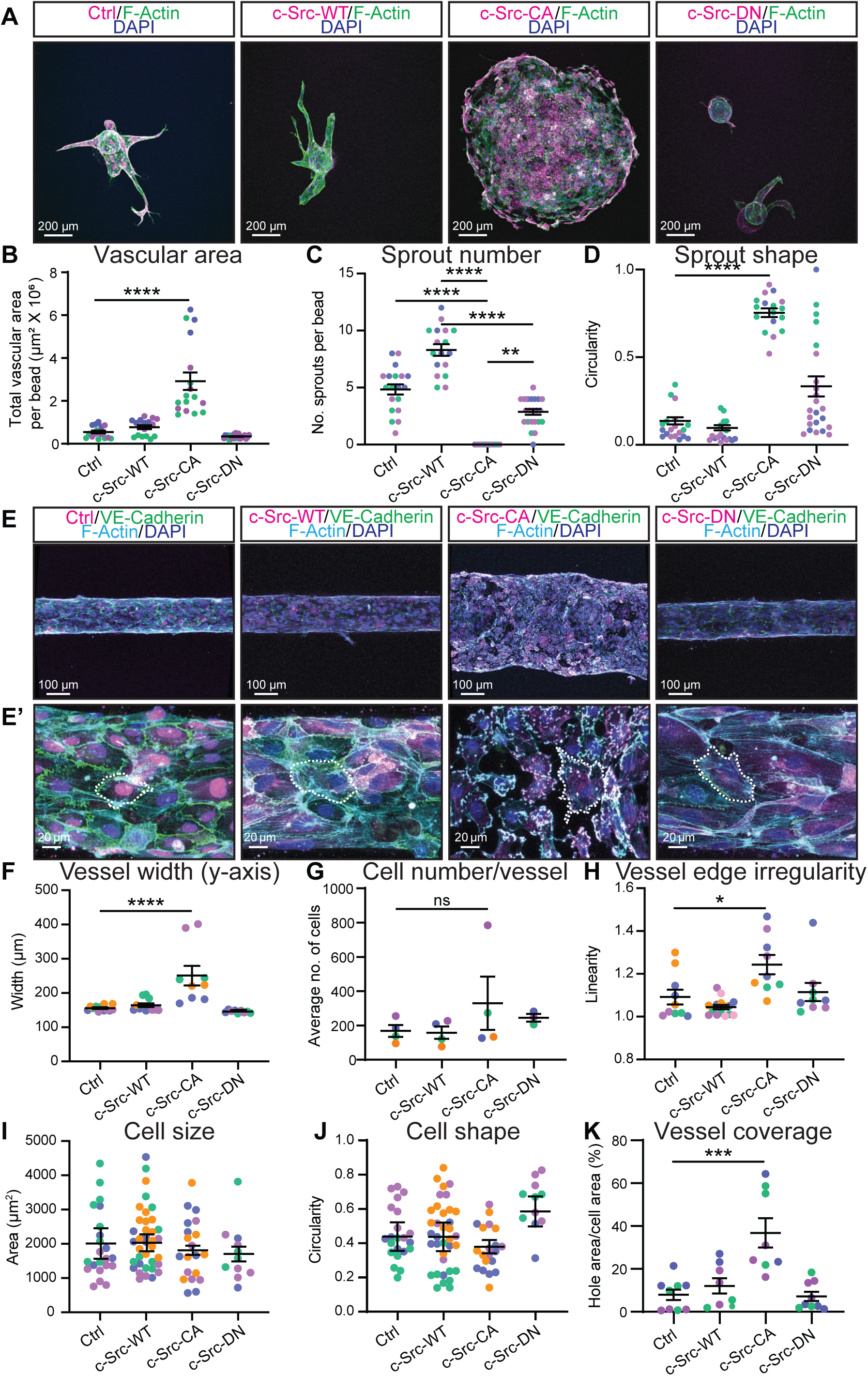
Constitutively active c-Src induces ballooning of *in vitro* 3D vasculature. **(A)** Representative images of fibrin bead sprouts. HUVECs transduced with mScarlet-tagged c-Src mutants (magenta) were grown in a 5 mg/mL fibrin gel bead sprouting assay for 7 days before fixation. Immunofluorescent staining was performed for F-Actin (Phalloidin; green) and nuclei (DAPI; blue). Quantification of sprout parameters of vascular area **(B)**, number of sprouts per bead **(C)**, and shape of sprouting area **(D)**. n=3 independent experiments. **(E)** Representative images of HUVECs transduced with mScarlet-tagged c-Src mutants (magenta) seeded in PDMS microfluidic vessels containing 2.5 mg/mL collagen matrix for 3 days before fixing. Immunofluorescent staining was performed for VE-Cadherin (green), F-Actin (cyan), and nuclei (DAPI; blue). Low magnification **(E)** was used to quantify vessel width **(F)**, number of cells per vessel **(G)**, and vessel edge irregularity **(H)** n = 3-5 independent experiments, 3 vessels per replicate with each data point representing an average of 3-5 measurements per vessel. High magnification **(E’)** was used to quantify individual cell morphology within microvessels as cell size **(I)**, cell shape **(J)**, and vessel coverage as a percentage of total vessel area **(K)** n = 3-5 independent experiments, 3-5 images per replicate with each data point representing an average of 1-5 cells per image. All data are represented as mean ± SEM with individual data point indicated and colours represent independent experiments. Statistical significance was determined using Kruskal-Wallis test with Dunn’s multiple comparisons. The mean value of each individual replicate and corresponding SEM, instead of all data points separately, was used for statistical analysis (I, J). * p<0.05, ** p<0.01, **** p<0.001.

To further investigate the role of c-Src activity in 3D, we assessed the behaviour of c-Src-mSc cells in microfabricated vessels [32, 33]. Here, ECs were surrounded by a type I collagen matrix and subjected to oscillatory flow (∼ >3 dynes/cm2) by rocking. Consistent with the observations in the bead sprouting assay, vessels comprised of c-Src-CA cells displayed a rapid ballooning morphology after 3 days of culture (Fig 1E, Supp Movie 2), resulting in a significant increase in vessel width compared to Ctrl cells (Fig 1F). Neither c-Src-WT or c-Src-DN microvessels displayed significant changes in vessel width, revealing that c-Src autoinhibition by Y527 phosphorylation is essential for preventing vascular ballooning. The ballooning of c-Src-CA was not due to alterations in cell proliferation as we did not observe significant changes in the number of cells per vessel (Fig 1G). This was further supported by our observation that there were no changes in proliferation in c-Src mutant cells grown in a 2D monolayer (Supp Fig 1F, H). Higher magnification analysis of the microvessels revealed that the c-Src-CA cells had an both an irregular perimeter and vessel edge (Fig 1E’, H), but no change in cell size or shape (Fig 1I, J). We found that a significant percentage of the c-Src-CA vessels lacked cell coverage resulting in gaps in monolayer (Fig 1K), however cells within the vessel still maintained some connections (Fig 1E’). Analysis in 2D culture revealed no change in cell size, while a significant increase in the cell perimeter and a loss of cell circularity in c-Src-CA cells was observed (Supp Fig 1G, I-K). This discrepancy in observed cell shapes might be due to morphological alterations in ECs when grown in 2D versus encapsulated 3D settings exposed to flow. Taken together, these results reveal that constitutive activation of c-Src in endothelial cells induces a malformed and perforated vasculature.

### Constitutively active c-Src induces phosphorylation of target proteins at focal adhesions and cell-cell junctions

A loss of c-Src in ECs leads to a reduction in cell-matrix adhesion [21], and loss of c-Src expression at cell-cell junctions results in reduced VE-cadherin internalisation [15, 34]. We hypothesised that increased c-Src activation would result in an induction of FA and/or VE-cadherin phosphorylation and internalisation. We observed a modest increase in FA size, number and density in c-Src-WT cells grown in 2D, which was further exacerbated by the CA mutation (Fig 2A-D). Like the 3D microvessels, c-Src-CA induced large gaps in the endothelial monolayer (Fig 2A), with punctate accumulations of VE-cadherin at sites of cell-cell contact. Furthermore, a significant increase in the phosphorylation of key FA components paxillin (at Y118) and FAK (at Y576) in c-Src-CA cells was observed in comparison to controls (Supp Fig 2A-C), in line with previous observations [35, 36]. FAK phosphorylation at Y397 was induced, which is known to be involved in the recruitment and activation of c-Src to FAs [37] (Supp Fig 2D-G). This suggests that c-Src acts both up and downstream of FAK. Phosphorylation levels of paxillin Y118 and FAK Y576/Y397 in c-Src-DN cells were comparable to Ctrl (Fig 2A-D, Supp Fig 2A-G), confirming that the K295 site is essential for the kinase activity of c-Src. Comparing phosphorylation of paxillin Y118 in 2D to the 3D microvessels revealed conservation of the increased FA size and density in c-Src-CA cells compared to Ctrl, WT and DN (Fig 2E-H).

**Figure 2.**
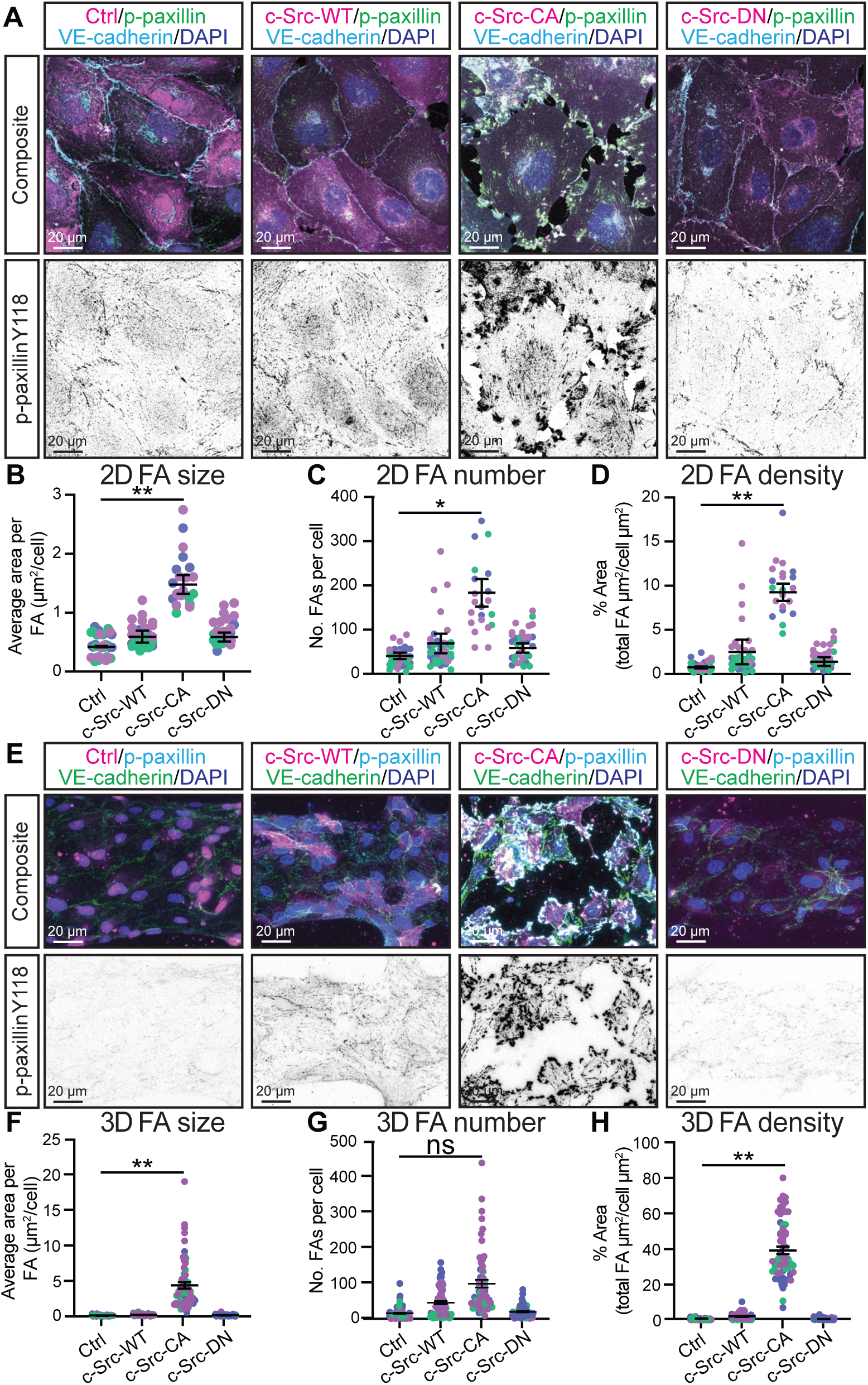
Constitutively active c-Src induces focal adhesion formation and disrupts cell-cell junctions in 2D and 3D. **(A)** Representative images of confluent monolayers of mScarlet-tagged c-Src mutant HUVECs (magenta), with immunofluorescent staining performed for focal adhesions (phospho-paxillin Tyr118; green, shown as black in individual panel), VE-cadherin (cyan) and nuclei (DAPI; blue). Quantification of average focal adhesion size **(B)**, number **(C)**, and density **(D)** per cell. n = 3 independent experiments, 5 images per replicate with each data point representing an average of 3-5 cells per image. **(E)** Representative images of HUVECs transduced with mScarlet-tagged c-Src mutants (magenta) seeded in PDMS microfluidic vessels containing 2.5 mg/mL collagen matrix for 3 days before fixing. Immunofluorescent staining was performed for focal adhesions (phospho-Paxillin Y118; cyan, shown as black in individual panel), VE-Cadherin (green), and nuclei (DAPI; blue). Quantification of average focal adhesion size **(F)**, number **(G)**, and density **(H)** per cell. n = 3 independent experiments, 4-5 images per replicate with each data point representing an average of 3-5 cells per image. All data are represented as mean ± SEM with individual data point indicated and colours represent independent experiments, and large circles represent average mean per replicate. Statistical significance was determined using Kruskal-Wallis test with Dunn’s multiple comparisons. The mean value of each individual replicate and corresponding SEM, instead of all data points separately, was used for statistical analysis (B-D, F-H). * p<0.05, ** p<0.01.

To investigate the cause of the loss of endothelial junction integrity (Fig 1E, K, and Fig 2A), we assessed the effect of elevated c-Src activity on VE-cadherin phosphorylation sites (Y568/Y731) previously reported to be linked to VEGFR2-dependent phosphorylation [3]. As expected, c-Src mutants did not alter VEGF-A-induced VEGFR2 phosphorylation at Y951 (Fig 3A, B), as this has been reported to be upstream of c-Src signalling [34]. Phosphorylation of VE-cadherin at Y658 or Y731 was not significantly altered by c-Src-WT or c-Src-DN compared to Ctrl, while c-Src-CA cells displayed a robust increase of VE-cadherin phosphorylation independently of VEGF-A stimulation (Fig 3A, C, D). Therefore, reduction of endothelial cell-cell contacts in c-Src-CA cells may be due to elevated VE-cadherin phosphorylation and subsequent internalisation.

**Figure 3.**
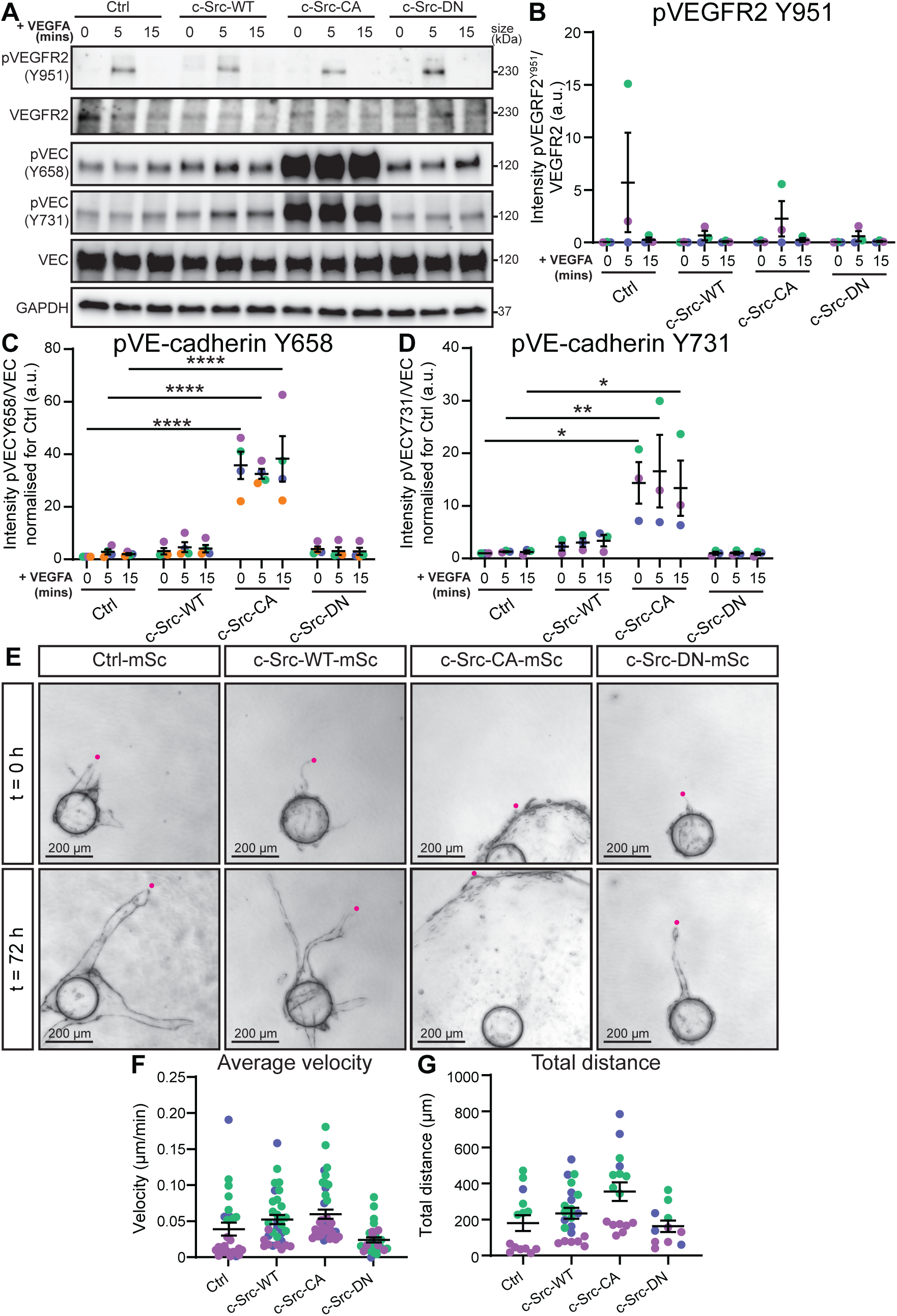
Constitutively active c-Src increased VE-cadherin phosphorylation and decreases 3D cell migration. **(A)** Representative images of Western blots of VEGFA-induced VEGFR2 and VE-cadherin activation in confluent monolayers of c-Src mutant HUVECs starved overnight in serum-reduced media before treating with 100 ng/mL VEGFA for 0 min, 5 min, or 15 min. Activation of VEGFR2, and VE-cadherin are shown by phosphorylation on indicated tyrosine sites. GAPDH was used as loading control. **(B)** Quantification of VEGFR2 phosphorylation at Tyr(Y)951 confirms receptor activation with a peak at 5 minutes. Ratio of phosphorylated VEGFR2 (Y951) to total VEGFR2 corrected to loading control (GAPDH). **(C)** Quantification of VE-cadherin phosphorylation at Y658 relative to total VE-cadherin corrected for loading control (GAPDH) and normalised to 0 min. **(D)** Quantification of VE-cadherin phosphorylation at Y731 relative to total VE-cadherin corrected for loading control (GAPDH) and normalised to 0 min. n = 3-4 independent experiments. **(E)** HUVECs transduced with mScarlet-tagged c-Src mutants (magenta) were grown in a 5 mg/mL fibrin gel bead sprouting assay for 5 days before live imaging at 30 min intervals over 72 h (representative images of 0 and 72 h). Quantification of total distance from the bead within 72 h **(F)**, or average velocity of the sprouting front **(G)**. n = 3 independent experiments. using Kruskal-Wallis test with Dunn’s multiple comparisons. All data are represented as mean ± SEM with individual data point indicated and colours represent independent experiments. Statistical significance was determined using 2way ANOVA with Tukey’s multiple comparisons (B-D) or Kruskal-Wallis test with Dunn’s multiple comparisons (F, G), * p<0.05, ** p<0.01, ***=p<0.005, ****=p<0.001.

We had previously identified that loss of FAs in endothelial cells results in a highly unstable vasculature due to the inability of cells to form functional contacts with their surrounding matrix [21]. However, while cells could not move in a co-ordinated manner, their velocity was unchanged. Thus, we next assessed the effect of c-Src mutants on EC migration in 2D and 3D. We found that c-Src-CA significantly reduced cell migration velocity and distance in 2D, whilst migration remained unaltered in cells expressing c-Src-WT or c-Src-DN (Supp Fig 3A- C, Supp Movie 3). The reduction in migration correlated with an increase in FA size c-Src-CA expressing cells. As reduction in cell-cell junction integrity has been shown to increase migratory capacity and sprouting angiogenesis [38], our data suggest that a balanced control of both cell-matrix and cell-cell junctions is essential for mediating migration. In contrast, when migratory capacity was assessed in 3D over three days (Fig 3E-G, Supp Movie 4), the average velocity of c-Src-CA cells was unchanged (Fig 3F) and the total distance travelled from the bead by c-Src-CA cells trended towards an increase, although this difference was not significant compared to Ctrl, c-Src-WT or c-Src-DN cells (Fig 3G). However, the differences in migration capacity of c-Src-CA cells observed in 2D (Supp Fig 3) and 3D (Fig 3) is likely due to the intricate challenges posed by migration through a mesh network of extracellular molecules, wherein cells exhibit the ability to move in a three-dimensional manner rather than adhering to linear movement patterns typically seen on matrix-coated glass surfaces [39].

### c-Src activation induces local degradation of the surrounding extracellular matrix

Our data demonstrated that c-Src-CA induced elevated paxillin (Y118), FAK (Y576) and VE-cadherin (Y658, Y731) phosphorylation and reduced endothelial cell-cell contacts. However, alterations in cell-matrix adhesion and cell-cell adhesion alone are not sufficient to explain the rapid expansion of c-Src-CA cells into the 3D fibrin (Fig 1A) or collagen (Fig 1E) matrix. FAs are localised sites of exocytosis in ECs [40], allowing secretion of proteases to subsequently remodel their surrounding environment in non-ECs [22]. Furthermore, unphosphorylated paxillin is known to be required for fibronectin fibrillogenesis [41]. Therefore, we hypothesised that upon elevation of c-Src activation, which significantly increases phospho paxillin Y118 and FAs number and size, fibronectin fibril assembly and ECM degradation may be altered.

We assessed the deposition of fibronectin fibrils over a time course ranging from 4-48 hours in c-Src mutant cells grown on uncoated glass coverslips. We found that c-Src-CA cells did not assemble fibrils over time (Supp Fig 4) in contrast to Ctrl, c-Src-WT and c-Src-DN cells which displayed elevated fibril assembly (48 vs. 4 hours). To ascertain if this was due to a reduction in fibronectin deposition or increased degradation, we grew ECs on fibronectin for 24 hours. This resulted in a significant decrease in fibril area underneath c-Src-CA cells which was rescued by the c-Src-DN mutation (Fig 4A-B). In addition to increased fibronectin degradation by c-Src-CA cells, we identified specific local fibronectin degradation at the sites of enlarged focal adhesions in c-Src-CA cells via Total Internal Reflection Microscopy (Fig 4C-D). These results suggested that FAs within ECs act as sites of local secretion of proteases to degrade surrounding ECM components.

**Figure 4.**
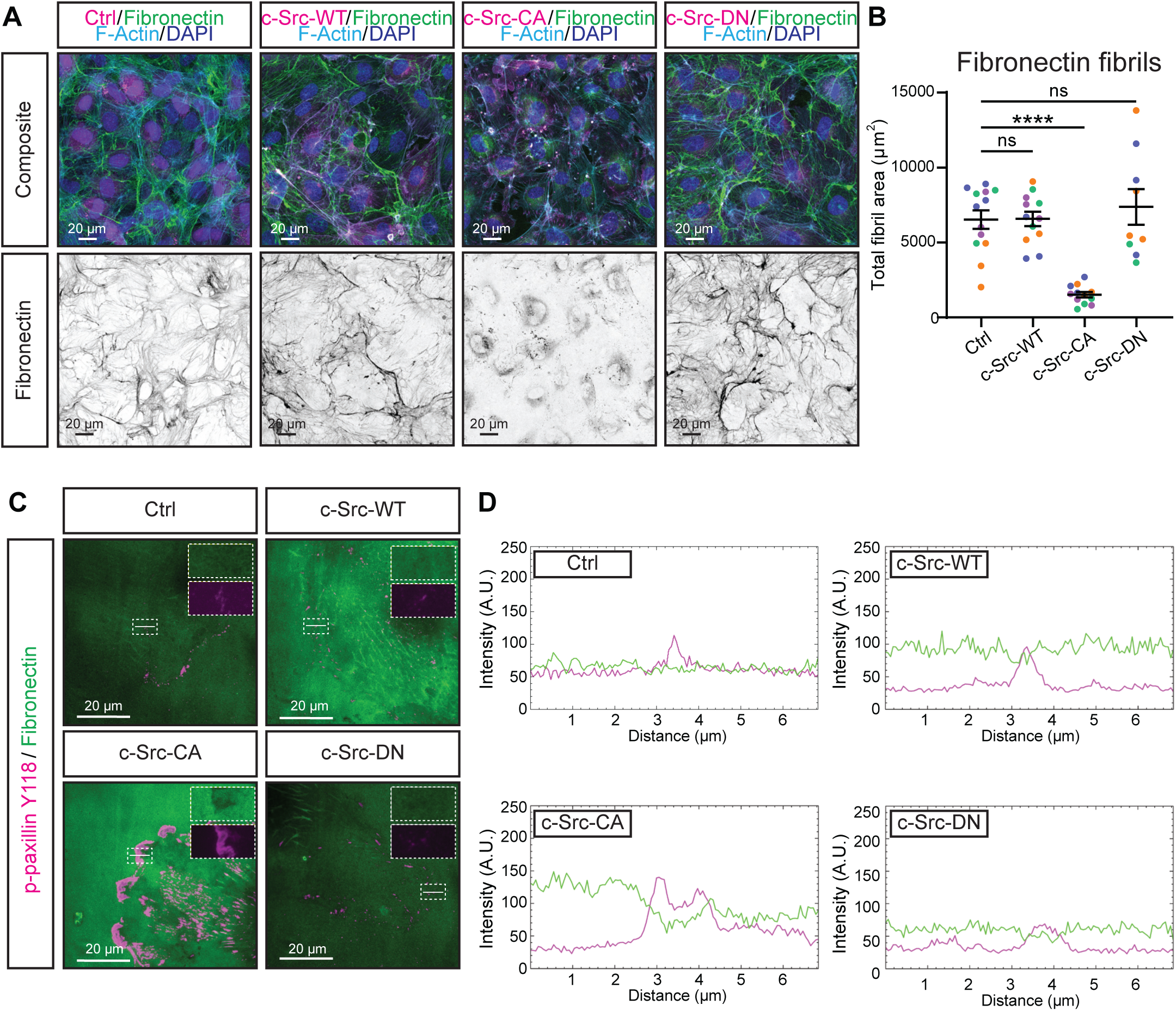
Constitutively active c-Src disrupts fibrillogenesis and degrades fibronectin locally at focal adhesions. **(A)** Representative images of mScarlet-tagged c-Src mutant HUVECs (magenta) grown on fibronectin-coated glass for 24 h before fixing. Immunofluorescent staining was performed for fibronectin (green, shown as black in individual panel), F-Actin (cyan), and nuclei (DAPI; blue). **(B)** Quantification of total fibronectin fibril area per image. n = 4 independent experiments, 3 images per replicate. Kruskal-Wallis test with Dunn’s multiple comparisons. **(C)** Representative images of total internal reflection fluorescence (TIRF) microscopy of sub- confluent mScarlet-tagged c-Src mutant HUVECs grown on fibronectin for 4 h before fixing. Immunofluorescent staining was performed for fibronectin (green) and focal adhesions (p- paxillin Y118; magenta). White dashed box indicates region of higher magnification show in top right corner, white lines indicate area used to generate intensity plots. **(D)** Intensity plots of areas of interest indicated by white lines in (C) to demonstrate a decrease in fibronectin (green) intensity at sites of increased focal adhesion signal (magenta). All lines are 7 µm and traverse the cell membrane through a focal adhesion. All data are represented as mean ± SEM with individual data point indicated and colours represent independent experiments. Statistical significance was determined using Kruskal-Wallis test with Dunn’s multiple comparisons. ****=p<0.001.

As proteases can be membrane inserted or secreted, we next addressed if the cells were releasing soluble proteases. We added conditioned media from Ctrl, c-Src-WT, c-Src-CA and c-Src-DN cells to wildtype ECs. As wildtype cells were still able to remodel fibronectin into fibrils upon addition of conditioned media obtained from c-Src-CA cells (Fig 5A-B), we concluded that the proteases secreted at FAs were not soluble, and were likely tethered to the cell membrane. This was further evidenced by mixing unlabelled wildtype ECs with mScarlet-labelled Ctrl, c-Src-WT, c-Src-CA and c-Src-DN cells in a bead sprouting assay. Ctrl, c-Src-WT and c-Src-DN cells formed sprouts containing both wildtype and mutant cells (Fig 5C-D). However, c-Src-CA cells exclusively formed balloons, while wildtype cells coated on the same bead maintained their ability to form sprouts. Taken together, these results reveal that proteases produced by c-Src-CA cells are locally secreted at FAs but are membrane bound.

**Figure 5.**
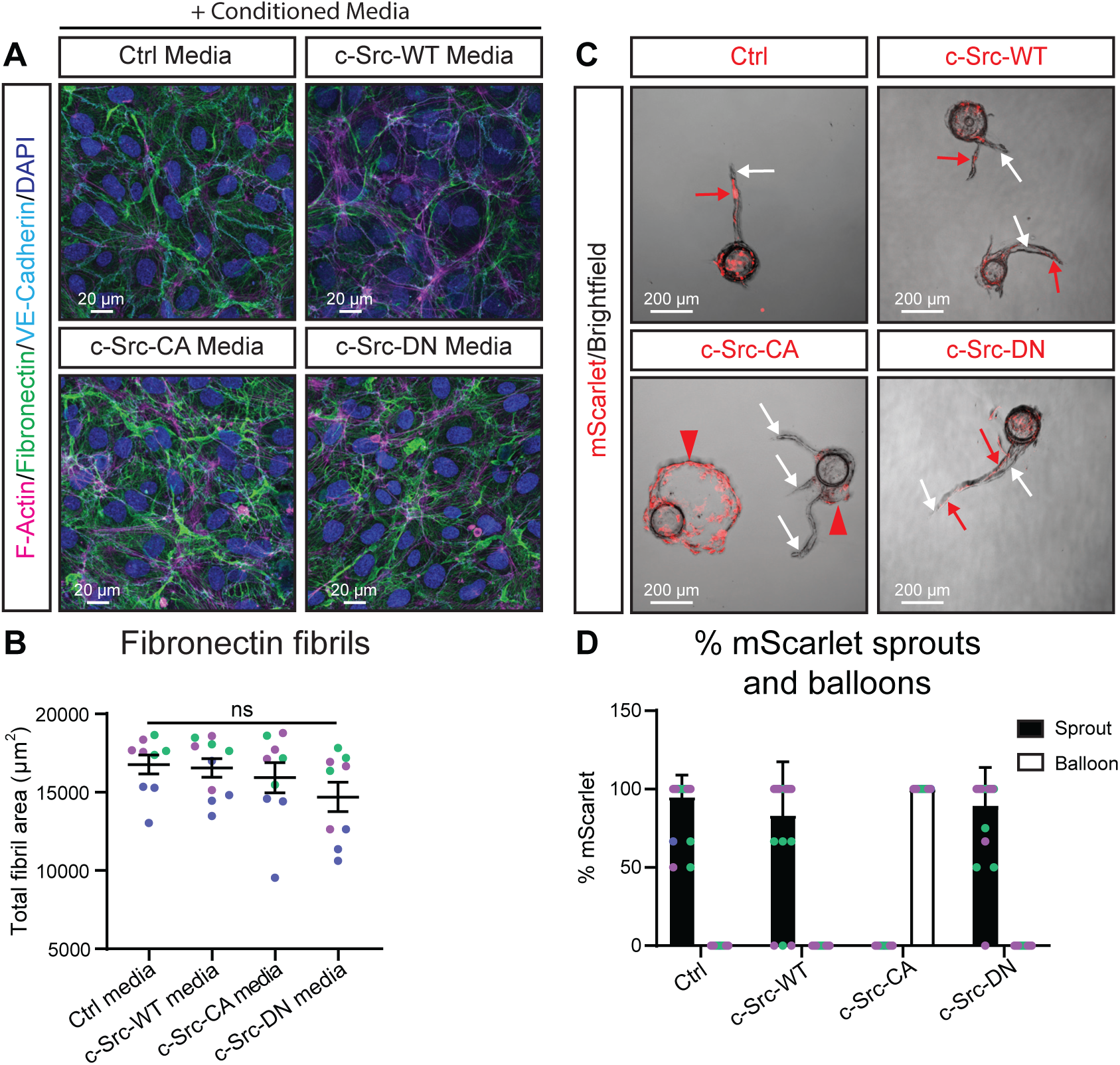
c-Src-dependent fibronectin degradation is mediated by insoluble factors. **(A)** Representative images of HUVECs grown on fibronectin-coated glass, treated with conditioned medium from c-Src mutant HUVECs. Cells were fixed and immunofluorescent staining was performed for F-Actin (magenta), fibronectin (green), VE-cadherin (cyan) and nuclei (DAPI; blue). **(B)** Quantification of total fibronectin fibril area per image upon treatment with conditioned medium from c-Src mutants. n = 3 independent experiments with 3 images per replicate. **(C)** Representative images of mixed bead sprouting assay of untransduced (brightfield) and mScarlet-tagged c-Src mutants transduced (red) HUVECs grown in a 5 mg/mL fibrin gel bead sprouting assay for 7 days before imaging. **(D)** Quantification of the percentage of sprouts or balloons positive for mScarlet cells. n = 3 for Ctrl and c-Src-CA, n = 2 for c-Src-WT and c-Src-DN. All data are represented as mean ± SEM with individual data point indicated and colours represent independent experiments. Statistical significance was determined using Kruskal-Wallis test with Dunn’s multiple comparisons.

### Inhibition of matrix metalloproteinases rescues vascular ballooning induced by constitutively active c-Src

MMPs degrade various proteins within the ECM and play a role in a range of vascular processes, including angiogenesis, morphogenesis and wound repair [26]. In non-endothelial cells, FAs have been shown to be sites of MMP secretion, mediating FA turnover and allowing for cell migration [25]. This suggests that excessive ECM breakdown in c-Src-CA cells by elevated MMP activity at FAs may result in rapid vascular ballooning. Addition of a broad-spectrum MMP inhibitor, Marimastat [42], to ECs grown in a 2D monolayer was able to rescue fibronectin degradation in c-Src-CA cells (Supp Fig 5A, B). However, c-Src-CA cells still maintained increased number and size of FAs compared to Ctrl, c-Src-WT or s-Src-DN cells (Supp Fig 5A, C-E). Treatment of ECs with Marimastat in a fibrin bead sprouting assay resulted in a rescue of the ballooning morphology observed in the c-Src-CA cells (Fig 6A-D). MMP inhibition also decreased the sprouting ability of Ctrl and c-Src-WT cells, with a decrease in the sprout number and increased circularity (due to decreased number of sprout protrusions) (Fig 6A, C, D).

**Figure 6.**
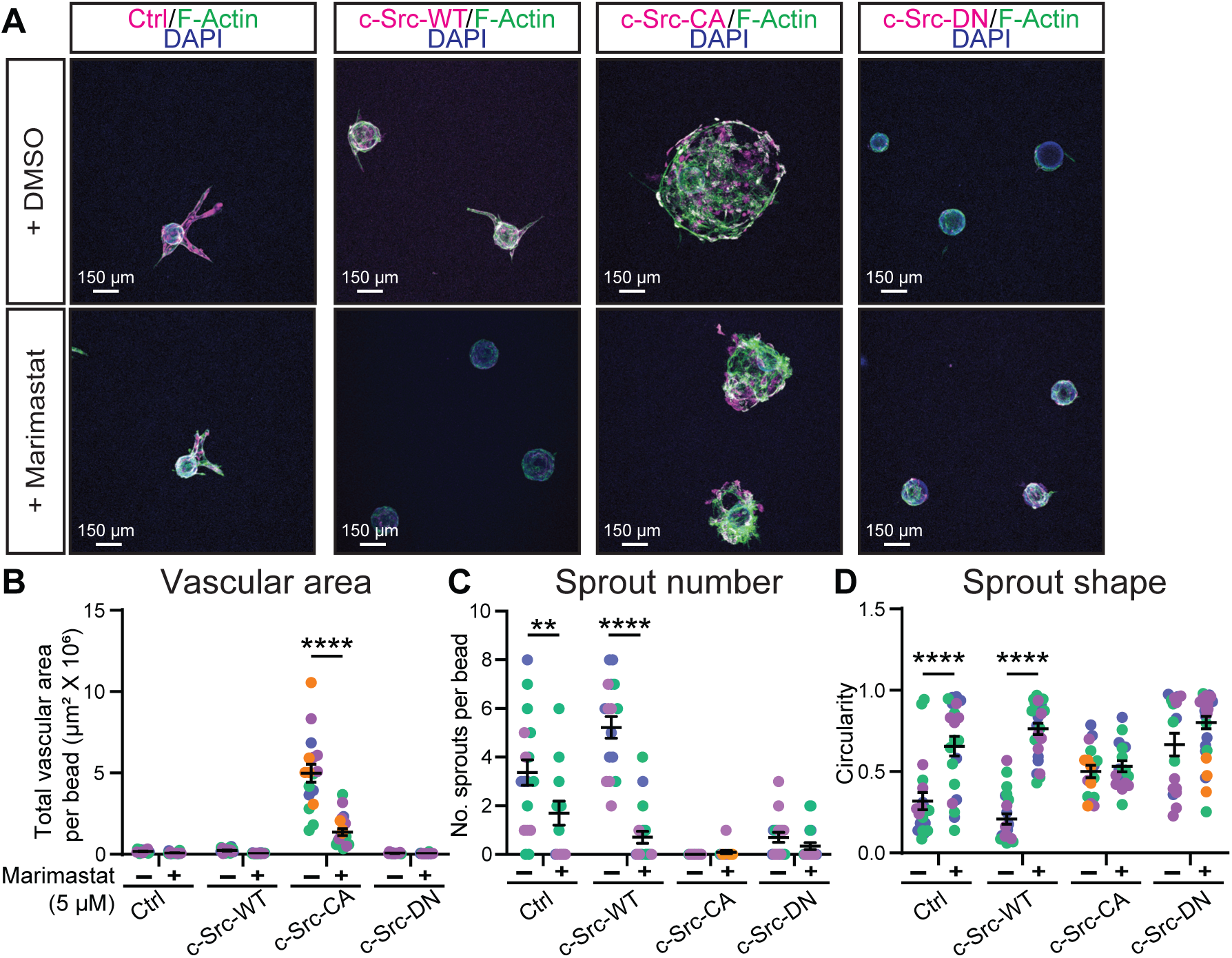
Broad spectrum MMP inhibitor Marimastat rescues vascular ballooning in constitutively active c-Src in 3D bead sprouting. **(A)** Representative images of mScarlet-tagged c-Src mutant HUVECs (magenta) grown in a 5 mg/mL fibrin gel bead sprouting assay for 7 days and treated every two days with DMSO vehicle control or 5 µM Marimastat before fixing. Immunofluorescent staining was performed for F-Actin (green) and nuclei (DAPI; blue). Quantification of sprout parameters of vascular area **(B)**, number of sprouts per bead **(C)**, and shape of sprouting area **(D)**. n=3 independent experiments. All data are represented as mean ± SEM with individual data point indicated and colours represent independent experiments. Statistical significance was determined using two- way ANOVA with Sidak’s multiple comparisons test. ** p<0.01, ****=p<0.001.

To investigate if MMP inhibition could also rescue the dysfunctional vasculature induced by c-Src-CA cells in a collagen matrix, 3D microvessels were treated with Marimastat. In Ctrl cells, Marimastat treatment had no effect on vessel width, vessel edge irregularity, vessel coverage, nor on cell size, shape and number (Fig 7A-G), suggesting that in a non-angiogenic setting, Marimastat does not adversely affect vessel morphology. In agreement with the results observed in fibrin bead sprouting (Fig 6), treatment of 3D microvessels with Marimastat rescued the increase in vessel width induced by c-Src-CA cells (Fig 7A, B). While large FAs were still detected in c-Src-CA cells treated with Marimastat (Supp Fig 5), the loss of c-Src-CA cell coverage of the vessel was restored (Fig 7A, G). Taken together, these results reveal that elevated c-Src activation induces large FAs, inducing enhanced local secretion of MMPs to degrade the ECM and induce vascular malformations.

**Figure 7.**
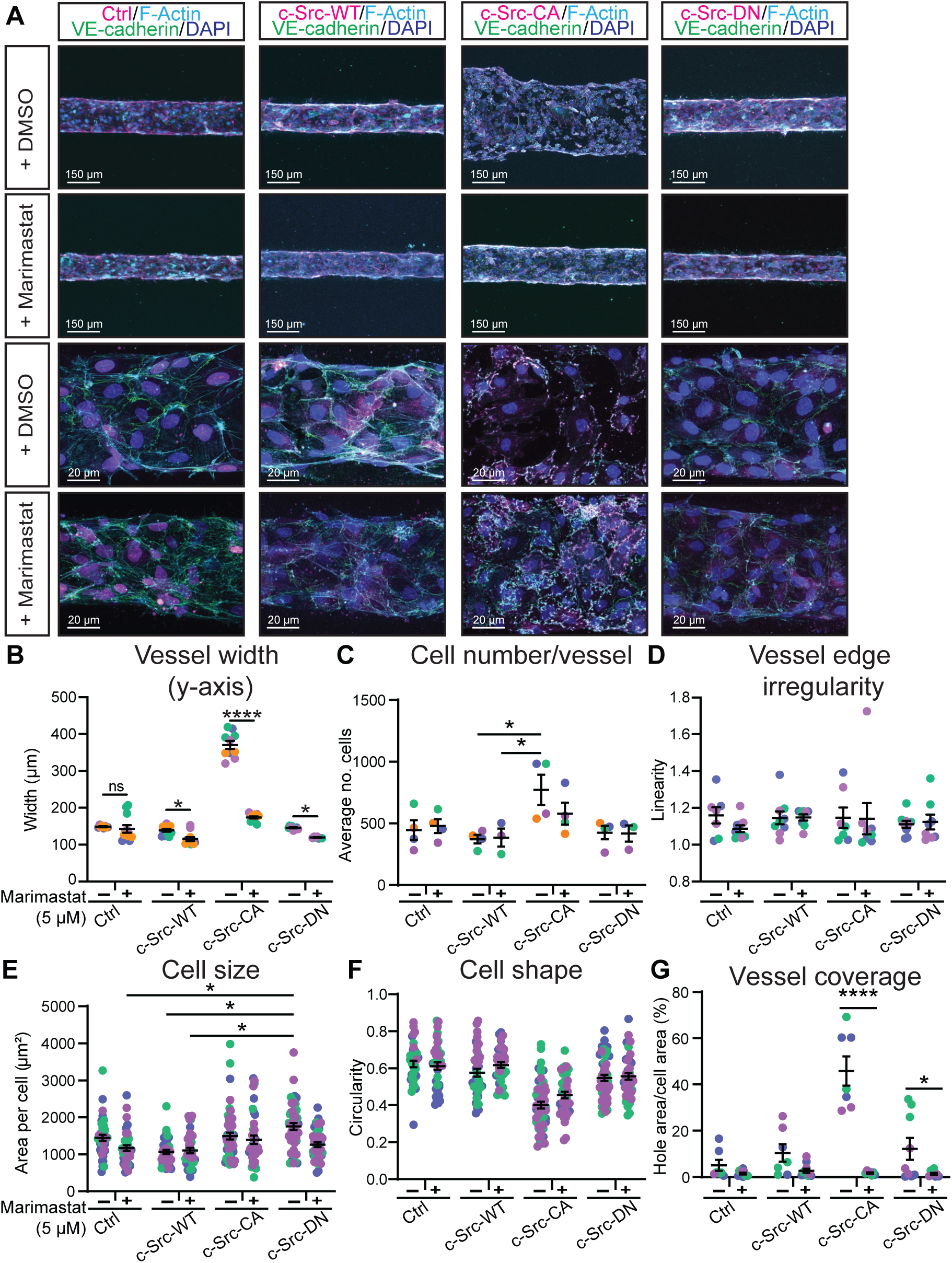
Broad spectrum MMP inhibitor Marimastat rescues vascular ballooning in constitutively active c-Src in 3D microvessels. **(A)** Representative images of HUVECs transduced with mScarlet-tagged c-Src mutants (magenta) seeded in PDMS microfluidic vessels containing 2.5 mg/mL collagen matrix for 3 days and treated daily with DMSO vehicle control or 5 µM Marimastat before fixing. Immunofluorescent staining was performed for VE-Cadherin (green), F-Actin (cyan), and nuclei (DAPI; blue). Low magnification (A top panel) was used to quantify vessel width **(B)**, number of cells per vessel **(C)**, and vessel edge irregularity **(D)**. n = 3-5 independent experiments, 3 vessels per replicate with each data point representing an average of 3-5 measurements per vessel. High magnification (A, bottom panel) was used to quantify individual cell morphology within microvessels as cell size **(E)**, cell shape **(F)**, and vessel coverage as a percentage of total vessel area **(G)** n = 3-5 independent experiments, 3-5 images per replicate with each data point representing an average of 1-5 cells per image. All data are represented as mean ± SEM with individual data point indicated and colours represent independent experiments. Statistical significance was determined using two-way ANOVA with Sidak’s multiple comparisons test. The mean value of each individual replicate and corresponding SEM, instead of all data points separately, was used for statistical analysis (E, F). * p<0.05, **** p<0.001.

## Discussion

Our investigation into the impact of a constitutively active variant of c-Src on endothelial dynamics has unveiled a pivotal role for c-Src in orchestrating FA formation, steering the course of MMP secretion and activity precisely at the nexus of cell-matrix adhesion. Here we show strategically confined MMP-mediated matrix degradation at FAs leads to a rapid and aberrant dilation of the vascular lumen within the context of 3D environments. However, this dilation can be effectively counteracted by administering a broad spectrum MMP inhibitor, Marimastat. These findings reveal a novel function of FAs as targeted loci of protease secretion within ECs, generating an environment which is conducive to promoting cellular migration in a complex 3D matrix.

The regulatory influence of SFKs on EC behaviour and vascular adhesion has been extensively investigated across diverse contexts. The existence of structural similarities among c-Src, Yes, and Fyn [43] coupled with the limited availability of tools for precise interrogation of the distinct roles of each SFK member within the endothelium, has presented considerable challenges. The recent adoption of conditional knockout mice with endothelial-specific modifications has yielded valuable insights into the distinct control exerted by individual SFK members over various signalling outputs and resultant biological phenomena. In the context of developing retinal tissue, we have found that loss of c-Src in ECs results in impaired cell-matrix adhesion without gross alterations in VE-cadherin localisation or phosphorylation [21]. In contrast, loss of Yes in ECs results in reduced VE-cadherin phosphorylation and endocytosis [18]. Here, we clearly demonstrate that induction of c-Src activity results in elevated VE-cadherin phosphorylation, an occurrence decoupled from growth factor instigation or induction of fluid forces. These findings conclusively affirm the capability of c-Src to phosphorylate VE-cadherin. It should be noted that the predominant SFK responsible for VE-cadherin phosphorylation is subject to contextual variability; c-Src appears to orchestrate phosphorylation and consequent permeability downstream of endothelial nitric oxide synthase (eNOS) activity [19], whereas Yes likely takes on a primary role in this process in response to fluid shear stress [18].

The controlled expansion of vascular lumens is primarily contributed to Rho GTPases- dependent regulation of actin contractility. Ras-interacting protein 1 (Rasip1) plays a role in enhancing the activity of Cdc42 and Rac1 while concurrently inhibiting RhoA [44]. Together with ArhGAP29, this can further block Rho associated kinase (ROCK) signalling, which normally acts to phosphorylate myosin light chain (MLC) and regulate F-actin contractility [44, 45]. The inhibition of either RhoA or ROCK leads to lumen expansion due to the disruption of actin contractility, while the absence of Rasip1 results in narrower lumens in developing vasculature [46, 47]. Interestingly, c-Src has been shown to disrupt RhoA signalling through p190 RhoGAP, leading to disruption of FAs and increased motility [48]. Additionally, c-Src can counteract ROCK signalling, promoting vessel stability [49]. This stands in contrast to the phenotypes observed here when c-Src is activated, namely large FAs and destabilised vasculature. This discrepancy may arise from differences in the extent of c-Src activation in these models, in agreement with requirement for a controlled balance in pMLC levels [50]. Phosphoproteomic screening for c-Src substrates in non-endothelial cells has identified numerous candidates associated with actin remodelling and adhesion, such as Filamin B, Tensin and p-130-Cas [36]. Therefore, c-Src clearly possesses the capability to promote actin-dependent processes, potentially inducing heightened tension at the membrane, which could lead to alterations in the linear distribution of VE-cadherin.

Whereas cell-matrix and cell-cell adhesions were traditionally considered distinct membrane components, it is now becoming clear that cell-cell junctions and FAs are intricately associated [5, 13, 14, 51, 52]. Significant crosstalk between both adhesion sites occurs via reciprocal binding to the actin cytoskeleton in addition to binding similar effector proteins like c-Src and FAK [52]. Disruption of FA components such as talin result in impaired stability of the vascular bed and altered VE-cadherin organisation at junctions [5, 6], whereas stabilisation of the cytoskeleton upon loss of talin restores VE-cadherin organisation and vascular stability, demonstrating the actin-mediated crosstalk between the two adhesion compartments [6]. As c-Src has been found to regulate both FAs and cell-cell junctions, it possesses the capacity to fine-tune EC migratory behaviour. Besides the direct interaction with FAs and cell-cell junctions, the above discussed effects and potential control of the actin cytoskeleton by c-Src would enable regulation of the crosstalk between the adhesion compartments. This adds an additional layer of complexity to the molecular mechanism of c-Src-mediated control of cell adhesion and migration. The precise mechanisms governing the assembly and disassembly of both FAs and cell-cell junctions by the constituents of these adhesion complexes, including c-Src, FAK, paxillin, talin, vinculin, and the actin cytoskeleton, represent a captivating area for future study.

In addition to mediating cell-matrix adhesion and crosstalk to cell-cell junctions, FAs have been reported to function as exocytosis hotspots to mediate local ECM degradation and FA detachment essential for migration [25, 41]. Our results confirm that increased FA size and number, due to increased phosphorylation of complex constituents, induces MMP-dependent ECM degradation. Active c-Src has been implicated in the co-ordination between the tubulin cytoskeleton and the apical membrane, directing polarised fusion events [53]. In non-endothelial cells, microtubules guide the trafficking of MMPs specifically to FAs in order to promote FA turnover. Our data suggests this phenomenon is conserved in the endothelium, where large FAs act as docking sites for microtubule-mediated MMP exocytosis, initiating local breakdown of the surrounding matrix.

In endothelial cells, c-Src is known to induce phosphorylation and expression of MT1-MMP [54], a membrane bound MMP which has been shown to enhance invasion and migration of ECs during angiogenesis [27, 55]. Therefore, it is likely that MT1-MMP trafficking to FAs is altered in our models. Interestingly, this is not the first instance of localised secretion occurring at sites of cell-matrix adhesion in the endothelium, as exemplified by the FA protein Zyxin, which is known to facilitate the secretion of von Willebrand factor (vWF) from Weibel-Palade bodies (WPBs) [56]. Zyxin mediates the dynamic reorganisation of actin filaments around WPBs at sites of exocytosis, promoting fusion and release of vWF. This illustrates that FAs act as microdomains for exocytosis, influencing various vascular processes. However, whether c-Src-induced exocytosis can similarly regulate the release of vWF and promote thrombi formation remains unknown.

Growing cells within a 3D network offers the advantage of applying mechanical forces, a crucial aspect when investigating processes influenced by mechanics, such as cell-matrix adhesion. Moreover, assessing cellular phenotypes within this 3D network permits the analysis of mechanical forces, including fluid flow, mechanical stiffness, and stretch [39, 57]. Notably, we consistently observed alterations in FAs across various 2D and 3D models, and the application of Marimastat effectively rescued matrix degradation, regardless of the environmental context. In our study, we observed similar phenotypes when cells were grown in collagen or fibrin, or with or without flow. Thus, in our models, vascular expansion appears to be independent of the specific extracellular matrix type surrounding the cells and is not contingent upon fluid flow. However, it is important to note that c-Src is a well-known mechanosensor [58] and can regulate how the endothelium responds to changes in fluid flow [16, 59–62], stiffness [7] and stretch [63–65]. Alterations in these mechanical forces could conceivably lead to changes in c-Src activity, potentially comparable to the levels observed in c-Src-CA cells, thereby triggering the induction of vascular malformations.

The interactions between the vasculature and the extracellular matrix are associated with a range of disease states. In the process of luminal expansion in vascular malformations and arteriovenous malformations (AVMs), vessels need to overcome the mechanical ECM barrier to allow for vascular remodelling and infiltration into a 3D space. Interestingly, pre-clinical models of hereditary haemorrhagic telangiectasia (HHT) exhibit elevated integrin and FA activity [66] in the first report linking endothelial cell-ECM interaction to vascular malformations. The requirement of elevated MMP secretion in diseases such as HHT is evidenced by the fact patients with AVMs show elevated MMP9 and MMP2 expression in plasma and tissue samples [67, 68]. Our data suggests that heightened c-Src activity causes MMP induced ECM degradation to enable expansion of vascular malformations. Whether c-Src activity is induced in HHT or in endothelial cells within AVMs remains to be determined. Notably, ECM degradation allows endothelial cells to create new blood vessels in stiff environments. Increased ECM stiffness, common in conditions like cancer, promotes angiogenesis and disrupts blood vessel barriers, elevating the risk of metastasis [27, 69, 70]. High levels of MMPs correlate with poor prognosis in cancer patients, and MMPs have been investigated for their involvement in mediating growth of the tumour vasculature [71, 72]. While c-Src expression is increased in human cancers and c-Src is a well-known promotor of tumour angiogenesis [73, 74], the specific mechanisms by which changes in ECM stiffness activate endothelial c-Src to induce MMP secretion remain unknown.

In conclusion, our research has yielded compelling evidence underscoring the importance of finely tuned regulation of cell-matrix interactions in the establishment of a robust and functional vascular system. Through our investigations, we have showcased that the signalling processes intrinsic to endothelial cells demand meticulous control. Specifically, our findings have highlighted the consequences of perturbations in c-Src activity levels, with both the loss of c-Src activity, as documented by Schimmel in 2020 [21], and the hyper-activation of c-Src as documented here, leading to the formation of aberrant vascular networks. This newfound understanding opens promising avenues for therapeutic intervention in diseases characterised by irregular vasculature. One such strategy revolves around the targeted modulation of endothelial-derived proteases, aimed at curtailing vascular invasion and uncontrolled expansion, which holds potential for ameliorating conditions associated with malformed vasculature.

## Acknowledgements

We thank Lena Claesson-Welsh, <u>Deb Barkauskas</u> and Nicholas Condon for helpful discussions. This work was performed in part at the Queensland Node of the Australian National Fabrication Facility. A company established under the National Collaborative Research Infrastructure Strategy to provide nano and microfabrication facilities for Australia’s Researchers. Microscopy was performed at the Institute for Molecular Bioscience Microscopy Facility which was established with the support of the Australian Cancer Research Foundation (ACRF) and incorporates the Dynamic Imaging, Cancer Biology Imaging and Cancer Ultrastructure and Function Facilities.

## Author Contributions

Conceptualisation: T.E., L.S., E.G.; Methodology and validation: T.E., L.S., T.Y., M.K., A.Y., I.N., S.S.; Formal analysis: T.E., L.S., A.Y.; Data curation: T.E., L.S., T.Y., M.K., A.Y., I.N. B.H.; Writing - original draft: T.E., L.S., E.G; Visualisation: T.E., L.S., E.G.; Supervision: A.S.Y., S.S., A.L., E.G.; Project administration: A.L., E.G. Funding acquisition: L.S., I.N., A.S.Y., S.S., A.L., E.G.

## Data and materials availability

All data are available in the main text or the supplementary materials.

## Materials and Methods

### Antibodies and dyes

The following antibodies were used: rabbit anti-phospho-c-Src (Tyr418) (Invitrogen, 44660G; ICC, 1:100), rabbit anti-phospho-VE-cadherin (Tyr658) (Invitrogen, 44-1144G, WB 1:1000), rabbit anti- phospho-VE-cadherin (Tyr731) (Invitrogen, 44-1145G, WB 1:500), rabbit anti-phospho-paxillin (Tyr118) (Invitrogen, 44-722G; WB, 1:1000, ICC, 1:200), mouse anti-c-Src GD11 (Millipore, 05-184; WB, 1:1000; ICC, 1:200), goat anti-VE-cadherin (R&D systems, AF938, WB 1:2000), goat anti-VEGFR2 (R&D systems, AF357, WB 1:1000), mouse anti-VE-cadherin-Alexa647 (BD Biosciences, 561567; ICC, 1:250), mouse anti-phospho-FAK (Tyr397) (BD Biosciences, 611806; ICC 1:200; WB, 1:1000), mouse anti-fibronectin (BD Biosciences, 610077; ICC, 1:200), rabbit anti-phosho-paxillin (Tyr118) (Abcam, AB4833; ICC, 1:100), mouse anti-MT1-MMP (Abcam, ab51074; ICC, 1:200), rabbit anti-Ki67 (Abcam, ab15580, ICC, 1:200), rabbit anti-paxillin (Cell Signaling, 2542, WB 1:1000), rabbit anti-phospho-FAK (Tyr576) (Cell Signaling, 3281; WB, 1:1000), rabbit anti- phospho-VEGFR2 (Tyr951) (Cell Signaling, 2471, WB 1:1000), rabbit anti-GAPDH (Cell Signaling, 2118; WB, 1:5000), mouse anti-FAK (Santa Cruz, sc-271126; WB, 1:1000), Phalloidin (Alexa Fluor 488 and 670, Cytoskeleton Jomar, PHDG1-A, PHDN1-A, ICC, 1:500).

Secondary antibodies conjugated with Alexa-488, Alexa-555 or Alexa-647 for immunofluorescent staining were obtained from Invitrogen and used at 1:400 for ICC. Horseradish peroxidase (HRP)-conjugated secondary antibodies were obtained from ThermoFisher Scientific and used at 1:5000 for Western blot.

### Cell culture

Human umbilical venous endothelial cells (HUVECs) from single donor (CC-2935 Lonza) were cultured in Endothelial Basal Media-Plus supplemented with SingleQuots Bullet Kit (EGM-Plus, CC-5035 Lonza) until passage 2-3 for 3D cultures and passage 3-7 for 2D cultures. Human lung fibroblasts (NHLF, CC-2512 Lonza) and Human Embryonic Kidney (HEK)-293T cells (12022001 CellBank) were cultured in Dulbecco’s Modified Eagle Medium (DMEM) with L-glutamine and sodium pyruvate (Invitrogen), containing 10% (v/v) heat-inactivated FBS and 100 U/mL penicillin and streptomycin (Life Technologies Australia). All cells were kept at 37°C and 5% CO2.

### Generation of c-Src mutants

pLV-CMV-IRIS-PURO-c-Src-mScarlet plasmid was used to the generation of Y527F and Y527F/K295R mutants via QuickChange II Site-Directed Mutagenesis Kit (Agilent Technologies, 200523) according to manufacturer’s protocol using the primers listed in Table 1.

**Table 1:**
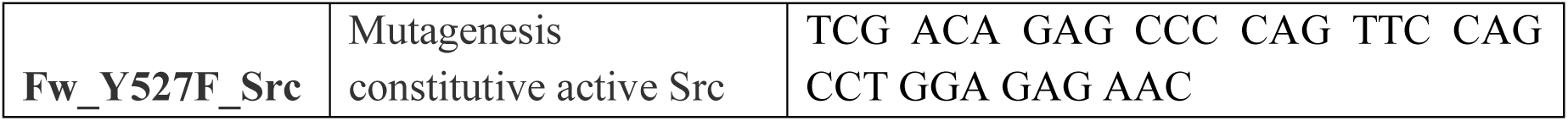
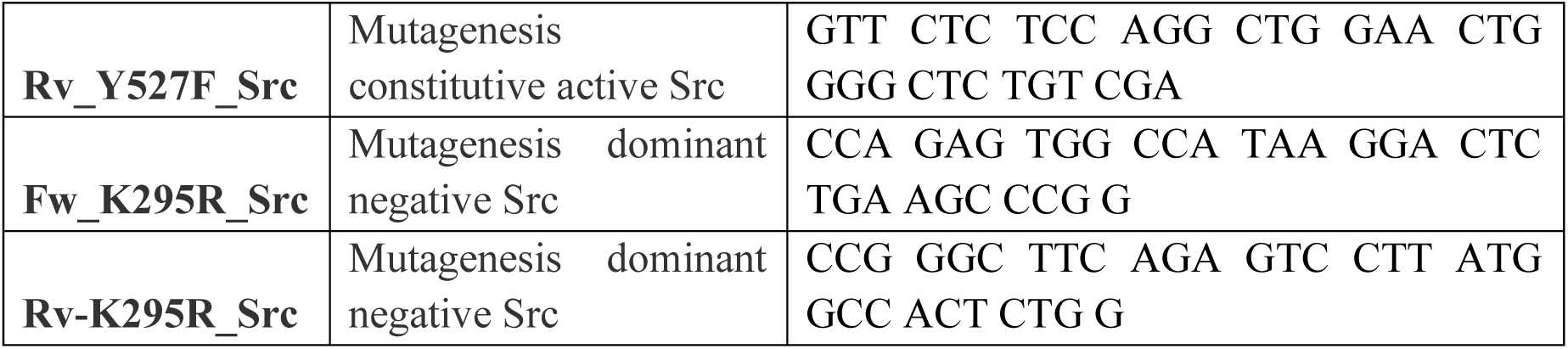
Cloning primers.

### Lentiviral transduction

Lentivirus constructs were packaged into lentiviral particles in HEK-293T cells by co-transfection of third-generation lentiviral packaging plasmids (pMDL/pRRE, pRSV-REV, and pMD2.G, kindly provided by Lena Claesson-Welsh, Uppsala University) with PEI 2500 (Polyplus) in OptiMEM (Gibco) according to manufacturer’s protocol. Supernatant containing lentivirus was harvested at 48 and 72 h after transfection, filtered through 0.45 µm filter and concentrated using Lenti-X (Scientifix, 631231). For lentiviral transduction, HUVEC were incubated with concentrated lentivirus, selected with puromycin (1.5 mg/mL) after 24 h and used for assays after 72 h.

### Bead sprouting assay

Fibrin gel bead sprouting assays were performed as described previously [31]. Briefly, HUVEC were coated onto Cytodex 3 microcarrier beads (Sigma-Aldrich) overnight. Coated microcarrier beads were embedded in a fibrin gel (fibrinogen (5 mg/mL, Sigma-Aldrich, F8630), aprotinin (50 µg/mL, Sigma-Aldrich, A3428) in EBM-Plus (Lonza, CC-5036) supplemented with 2% fetal bovine serum (Thermo Fisher Scientific, 10099141). Fibrinogen solution containing the microcarrier beads was clotted using 1 U of thrombin (Sigma-Aldrich, T9549/T4393) for 30 min at 37°C in a 24 well ibiTreat U-plate (Ibidi, 82406). Fibroblasts (NHLF) (20,000 cells per well) were added on top of the fibrin gel in EGM-Plus with 100 ng/mL VEGF-A (Thermo Fischer Scientific, PHD9391) and medium refresh every other day.

Live imaging was performed between days 3-7. Z-stacks were acquired every 30 min for 72 h at 37°C and 5% CO2 using Zeiss Axio Observer 7 inverted widefield microscope.

After 7 days sprouts were fixed and stained for ICC. Fibroblasts were removed using Trypsin and gels were thoroughly washed with PBS^+/+^ (PBS supplemented with 1 mM CaCl2 and 0.5 mM MgCl2) before fixing with 4% PFA for 10 min, permeabilising with 0.5% TritonX100 for 5 min, and blocking with 5% BSA in PBS^+/+^ for 2 h at room temperature on a shaker. Primary and secondary antibodies were incubated overnight at 4°C in 5% BSA in PBS^+/+^. Sprouts were imaged with a Zeiss LSM 710 Confocal microscope with Plan Apo 10x/0.45 objective.

### Microfluidic devices

Microfluidic devices were prepared as described previously [32] using 2.5 mg/mL Collagen I (rat tail, R&D Systems) as extracellular matrix. Channels were seeded with HUVEC at 1 x 10^6^ cells per mL and maintained under oscillatory, gravity-driven flow by lab rocker (>3 dynes/cm^2^) for 3 days at 37°C with 5% CO2. Media was refreshed daily.

Microvessels were fixed subsequently with 1% PFA containing 0.05% TritonX-100 for 90 sec, followed by 4% PFA for 15 min, and permeabilised in 0.5% TritonX-100 for 15 min all at 37°C. Microvessels were blocked in 2% BSA in PBS^+/+^ for 4 h at room temperature. Primary antibodies were incubated overnight at 4°C in 2% BSA in PBS^+/+^. Microvessels Z-stack images were acquired on a Zeiss LSM 880 Confocal microscope with 10x/0.45 Air or 40x/1.2 Water immersion objectives.

### Western blot (WB)

Cells were washed once with PBS^+/+^ and lysed in SDS-sample buffer containing 100 mM DTT and boiled at 95°C for 10 min to denature proteins. Proteins were separated by SDS-PAGE on 4-15% gradient gels (Mini-PROTEAN Precast gels, Bio-Rad) in running buffer (200 mM glycine, 25 mM Tris, 0.1% SDS (pH8.6)) and transferred to nitrocellulose membrane (Bio-Rad, 1620112) in blot buffer (48 nM Tris, 39 nM glycine, 0.04% SDS, 20% MeOH). Membranes were blocked in 5% (w/v) BSA (Sigma) in Tris-buffered saline with Tween 20 (TBST) for 30 min before incubating in primary antibodies overnight at 4°C followed by secondary antibodies linked to HRP (Invitrogen) for 1 h at room temperature. Between each step, the membranes were washed 3 times for 10 min in TBST. The HRP signals were visualised by enhanced chemiluminescence (ECL) (Bio-Rad) and imaged with a Chemidoc (Bio-Rad). Images were analysed using FIJI, measuring signal intensity, and adjusted to the loading control. Phosphorylated proteins were adjusted to the relative total protein.

### Immunocytochemistry (ICC)

For 2D immunofluorescent staining, HUVECs were cultured on 12 mm glass coverslips, or in 35 mm glass bottom dish (MatTek, P35G-1.5-14-C), coated with 5 µg/mL fibronectin (Sigma). Cells were washed with PBS^+/+^, fixed in 4% PFA for 10 min, and blocked/permeabilised for 30 min in 3% BSA, 0.3% TritonX-100 in PBS^+/+^. Primary antibodies were incubated for 60 min at room temperature in 1.5% BSA in PBS^+/+^, washed 3x in PBS^+/+^, and incubated in secondary antibody for 60 min at room temperature in 1.5% BSA in PBS^+/+^ before mounting in ProlongGold +DAPI (Cell Signaling Technologies). Imaging was performed on Zeiss LSM 710 Confocal microscope with 10x/0.45 Air, 40x/1.1 Water immersion, or 63x/1.15 Water immersion objectives. Or on Zeiss LSM 880 Confocal microscope with 10x/0.45 Air, 40x/1.2 Water immersion, 40x/1.3 Oil immersion, 63x/1.4 Oil immersion objectives.

### Conditioned Media

Puromycin selected HUVEC transduced with control or c-Src mutant lentivirus were grown on fibronectin coated glass coverslips and supernatant was harvested after 24 h. The conditioned medium was transferred onto wildtype HUVECs grown on fibronectin coated glass coverslips and incubated for 24 h before fixation and ICC staining with fibronectin and VE-cadherin.

### Drug Treatments

HUVECs were treated with 5 µM Marimastat (Sigma-Aldrich, M2699, in DMSO) or 0.01% DMSO as vehicle control in EGM-Plus medium. In 2D experiments, HUVECs were adhered for 2-4 h before Marimastat or control was administered for 24 h. Similarly, in microvessels, HUVECs were adhered for 2-4 h before Marimastat treatment once per day for 3 days. In fibrin sprouting assays, HUVECs were coated on beads and embedded in the fibrin gel for 24 h before Marimastat treatment was administered on days 1, 3, and 5, before fixation at day 7. For VEGF- A stimulation, confluent HUVECs were starved overnight in EBM-Plus with 1% FBS prior to addition of 100ng/mL VEGF-A165 (Thermo Fisher Scientific, PHC9391, in 0.1% BSA/PBS) in starvation medium.

### TIRF microscopy

HUVECs were grown for 4-6 h at sub-confluency on #1.5 coverslips (Hurst Scientific, 0117500) before fixing and staining as regular ICC. Critical angle was set for each condition using Nikon Ti2 Inverted Microscope Stand with Andor Dragonfly Spinning Disc Scanhead with 100x/1.49 TIRF Oil objective.

### Migration assay

HUVECs transduced with control-mScarlet or c-Src-mScarlet mutants and puromycin selected were seeded into glass bottom 24-well plate (Ibidi, 82426) containing a silicon barrier (Ibidi, 80209) and grown until confluent. The silicon barrier was removed, and time-lapse imaging of the scratch area was acquired for 16 h with 10 min interval on Nikon Deconvolution Ti-E Inverted Microscope with 10x/0.45 Air objective.

### Image Analysis and Quantification

For focal adhesion analysis in FIJI, maximum Z-projections were made, and filtered using Gaussian Blur (sigma=1) before sharpening. The focal adhesion signal was thresholded using the MaxEntropy, individual cells were traced using VE-cadherin border, and focal adhesion count, area, and density were measured within each cell using particle analyser (size: 0.25 µm^2^ - infinity).

Bead sprouting area was quantified in FIJI by tracing the perimeter of the sprouting area in a minimum Z-projection of brightfield image and subtracting the area of the bead. Number of sprouts was measured as number of lumenised vessels extending from the bead. Sprout length was measured from the edge of the bead down the centre of each sprout to its longest tip or as the greatest distance from the edge of the bead to the sprouting front of the cells for cells that did not sprout. Only sprouts that were growing horizontal through the gel were measured.

Microfluidic vessel width was measured in two axes (Y and Z) in Imaris. Five evenly spaced points across the tube were measured with a straight line and the mean was calculated per image to produce 2-3 length measurements per vessel. Linearity was measured by tracing the edge of the vessel (marked by phalloidin) and dividing this length by the length of a straight line through the vessel edge in FIJI. To measure the number of cells per image, Imaris spot counter was used to identify and quantify DAPI.

Total internal reflection fluorescence (TIRF) critical angle was adjusted in accordance with variations in coverslip thickness. Images were used to analyse intensity plots in FIJI, which were generated with a line of 7 µm length and 3 µm width. The focal adhesion channel was processed by background subtraction (rolling ball radius = 50.0), followed by Gaussian blur (sigma = 1.0).

Proliferation was quantified in sub-confluent cells by counting number of nuclei containing Ki- 67 as a ratio to total number of nuclei stained with DAPI.

Fibrillar area was calculated by subtracting background (rolling ball radius = 50.0) and thresholding the fibronectin channel using MaxEntropy and measuring area.

